# GRIPT: A novel case-control analysis method for Mendelian disease gene discovery

**DOI:** 10.1101/454975

**Authors:** Jun Wang, Li Zhao, Xia Wang, Yong Chen, Mingchu Xu, Zachry T. Soens, Zhongqi Ge, Peter Ronghan Wang, Fei Wang, Rui Chen

**Affiliations:** Human Genome Sequencing Center, Baylor College of Medicine, Houston, TX 77030, USA; Department of Molecular and Human Genetics, Baylor College of Medicine, Houston, TX 77030, USA; Structural and Computational Biology & Molecular Biophysics Graduate Program, Baylor College of Medicine, Houston, TX 77030, USA; Baylor Miraca Genetics Laboratories, Houston 77030, TX, USA; Shanghai Key Lab of Intelligent Information Processing, School of Computer Science and Technology, Fudan University, Shanghai, China

**Keywords:** Mendelian disease, disease gene prioritization, cohort analysis, locus heterogeneity, Next generation sequencing

## Abstract

Despite rapid progress of next-generation sequencing (NGS) technologies, the disease-causing genes underpinning about 50% of Mendelian diseases remain elusive. One main challenge is the high genetic heterogeneity of Mendelian diseases in which similar phenotypes are caused by different genes and each gene only accounts for a small proportion of the patients. To overcome this gap, we developed a novel method, the Gene Ranking, Identification and Prediction Tool (GRIPT), for performing case-control analysis of NGS data. Analyses of simulated and real datasets show that GRIPT is well-powered for disease gene discovery, especially for diseases with high locus heterogeneity.

## Background

Mendelian diseases refer to the diseases caused by mutations in a single gene and are inherited following Mendel’s laws. It was estimated that approximately 0.4% of live-born individuals have clinically recognizable Mendelian phenotypes by early adulthood, and about eight million children worldwide are born each year with a serious genetic condition leading to disability or threatening lives [1, 2]. Identification of Mendelian disease-causing genes can directly improve molecular diagnosis and genetic counseling and also provide new insights into the genetic and pathogenic mechanisms underlying the diseases, laying the foundations for developing preventive and therapeutic methods for patients [3, 4].

Traditional strategies for Mendelian disease gene discovery are primarily family-based approaches. Linkage analysis was widely used for mapping genes underlying dominant inherited diseases, while homozygosity mapping was successfully applied on recessive inherited diseases in consanguineous families [5-9]. However, family-based strategies are limited by the availability of multi-member families and cannot be effectively applied to the sporadic cases of rare diseases. On the other hand, as the recent advances in next-generation sequencing (NGS) technology and the establishment of large patient cohorts, case-control analysis of patient NGS data has provided powerful alternatives in novel disease gene discovery [7, 10]. Case-control analysis methods typically map candidate genes mutated in multiple affected patients (i.e. cases) but in wildtype form in unaffected individuals (i.e. controls). However, it remains challenging for these methods to distinguish the candidate disease genes from the genes with large numbers of rare benign variants (e.g. the highly mutable genes). Furthermore, the enormous amount of data generated by NGS brings huge analytical and computational burdens, which requires algorithms that can efficiently search through large numbers of whole genome/exome data and reliably detect the true signal of the disease gene from the massive background noise.

Previously, for case-control analysis, association tests were developed to identify the relation between genotypes and the phenotype, such as rare variant vs. common complex diseases. Particularly, the group-wise (i.e. gene/locus-based) association tests have been applied to enrich association signals and reduce the penalty for multiple testing. For example, “burden tests” or “collapsing methods”, such as Combined Multivariate and Collapsing (CMC) [11], Cohort Allelic Sums Test (CAST) [12], and Weighted-Sum method [13] aggregate prioritization information across multiple variants within a genetic region. Furthermore, the kernel-based methods, such as Sequence Kernel Association Test (SKAT)[14] and Kernel-Based Adaptive Clustering (KBAC)[15], take into account the different effect direction and magnitude of variants within a locus when grouping the variants together. However, these methods were not originally designed for Mendelian diseases. Moreover, most of these methods are mainly based on the allele frequency differences and take little account of the functional predictions of individual alleles. In 2011, a case-control analysis method named Variant Annotation, Analysis and Search Tool (VAAST) and, later, an upgraded version, VAAST2, were developed for disease gene discovery of Mendelian disorders [16, 17]. VAAST/VAAST2 measures the aggregative impact of variants within a gene based on the variant frequency differences between cases and controls, and also considers the functional effects of variants by weighting amino acid substitution frequency and phylogenetic conservation [16, 17]. However, VAAST/VAAST2 is prone to producing false positives, prioritizing the genes with large numbers of rare benign variants as the candidate disease genes. In addition, its specificity is greatly reduced when analyzing cohorts with high population stratification.

So far, 3532 genes underlying 5159 Mendelian phenotypes have been discovered, according to the Online Mendelian Inheritance in Man (OMIM) database (OMIM statistics, May 11th, 2018) [18]. But the genes mutated in about 50% of the known Mendelian disorders remain elusive, and many more Mendelian phenotypes have not yet been recognized [10]. One main challenge is that the disease is often rare and genetically heterogeneous where each disease-causing gene only accounts for a very small proportion of patients with the disease [10]. To address this challenge, we developed a novel method, named the Gene Ranking, Identification and Prediction Tool (GRIPT), to identify Mendelian disease genes through analyzing genomic sequence data of patient-control datasets. By testing both simulated and real datasets, we demonstrated that GRIPT has excellent sensitivity and specificity in identifying known and novel disease genes. It significantly outperforms other state-of-the-art tools in discovering disease genes underlying patient cohorts with high locus heterogeneity. Moreover, GRIPT is quite robust and less affected by potentially confounding factors, such as patient cohort size, population stratification in cohorts, and cutoff of variant frequency filtering.

## Results

### The framework of GRIPT

GRIPT is specifically designed for Mendelian disease gene discovery through prioritizing genes with significantly higher deleterious mutation load in patients than controls as the candidate genes. In implementation, GRIPT first ranks the variants within each gene for every individual in both patient and control cohorts according to the variant effect score provided by users, e.g. CADD score (Figure 1, see Methods). Based on the variant scores, a gene score is calculated for each gene measuring the deleterious mutation load of the gene in every individual under a given inheritance model, i.e. Autosomal dominant (AD), Autosomal recessive (AR), X-linked dominant (XD), or, X-linked recessive (XR) model (see Methods). Then, a Fisher’s test built upon the combination of a binomial test and a Wilcoxon rank sum test (WRST) is applied to compare the gene score distributions in patients and controls for each gene, and a significance p-value associated with the test statistic is assigned. This composite test is especially suitable to compare two highly skewed distributions with excesses of zero, such as the gene score distributions in the case and control cohorts (Figure 2, see method) [19]. Finally, GRIPT compares and ranks all genes based on the test statistic of each gene (Figure 1).

**Figure 1.**
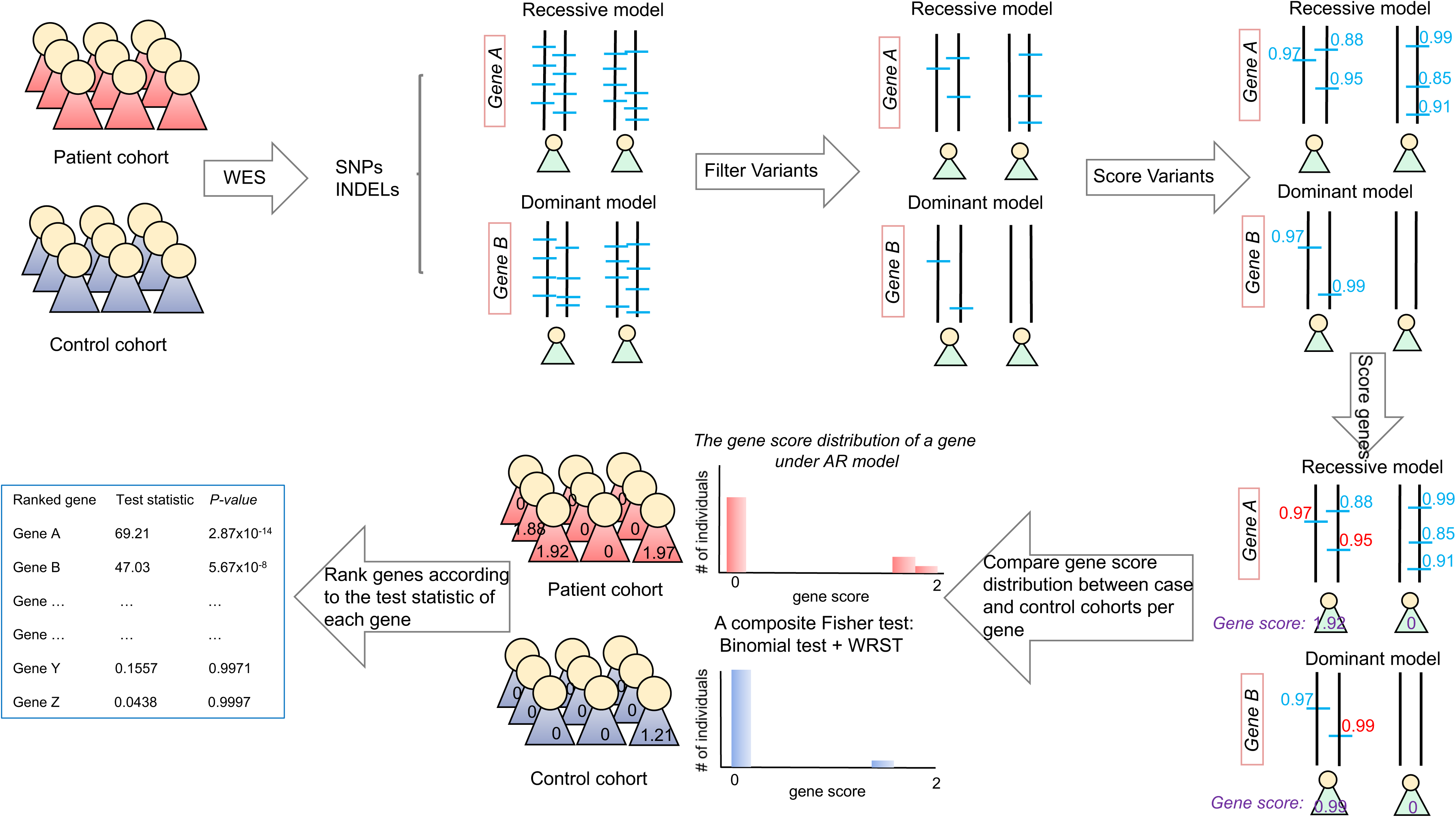
The logic flowchart of GRIPT. First, the samples of the case and control cohorts will be collected and be subjected to NGS, e.g. WES. After variant calling, the known common and/or benign variants will be filtered out based on the variant annotation and their allele frequency in large databases of normal populations. Thus, for each gene, only a few rare variants will be left. Then GRIPT will annotate and rank the deleteriousness of each variant, e.g. using CADD score. Based on the variant scores, a gene score will be calculated to measure the deleterious mutation load of each gene in every individual according to a given inheritance model (see Methods). Next, a Fisher’s test built upon the combination of a binomial test and a Wilcoxon rank sum test (WRST) will be calculated to measure the difference of gene score distributions between patient cohort and control cohort for each gene, and a significance p-value associated with the test statistic will be assigned. This composite test is especially well suited to measure the difference of two highly skewed distributions with excesses of 0, such as the gene score distribution in the patient/control cohort computed by GRIPT (Figure 2). Finally, according to the test statistic of each gene, GRIPT compares and ranks all genes.

**Figure 2.**
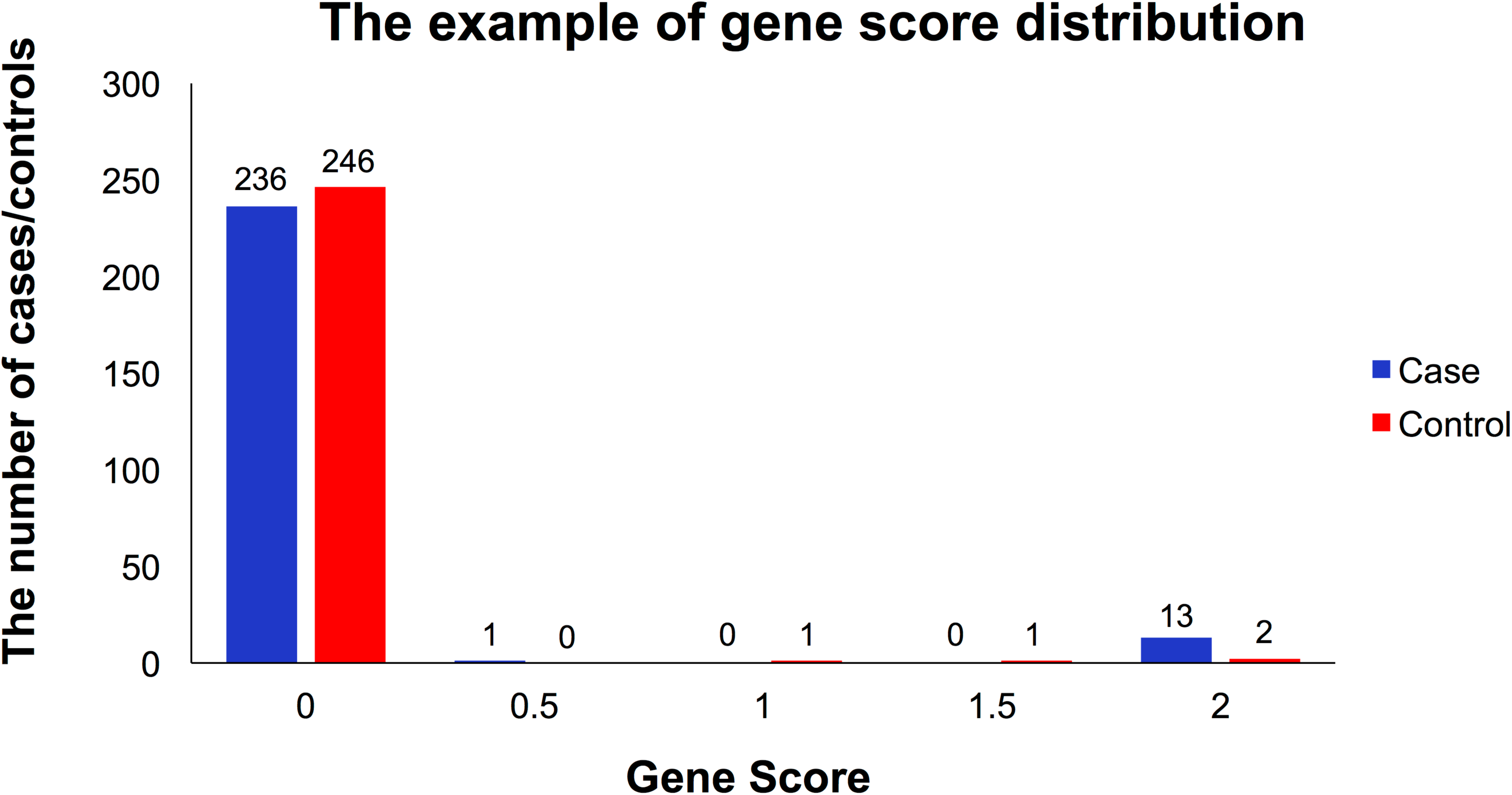
The example of gene score distribution. This figure shows the gene score distributions of *USH2A* in a retinal disease cohort of 250 patients (in red) and in a control cohort of 250 individuals (in blue). X axis: the gene score of *USH2A* per individual. Y axis: The numbers of patients or controls with the corresponding score. Like the gene *USH2A*, the gene score distributions of most genes are highly skewed with excesses of zeros.

### Simulation analysis tests the sensitivity and specificity of GRIPT

To evaluate the sensitivity and specificity of GRIPT, we simulated WES data for patient and control cohorts under both the AR and AD inheritance models based on the variant profile of the human genome in the ExAC database (see Methods). To mimic the patient cohort with high disease-locus heterogeneity where a given disease gene only accounts for a small proportion of the patients, pathogenic mutations of the same gene was randomly selected from The Human Gene Mutation Database (HGMD) and spiked into a small proportion (e.g. 0.5%, 1%, 2%, or 3%, respectively) of individuals in the patient cohort (see methods). The size of patient cohort was set at 600 and the control cohort at 5000. The simulation for each scenario was repeated 30 times. A genome-wide statistical significance level (GWSL) of 2.7×10^-6^ was used as the significant p-value cutoff for multi-testing correction (given about 18500 autosomal protein-coding genes annotated by RefSeq genes). The performance of GRIPT was measured with three parameters: 1) the ranking of the disease gene with spike-in pathogenic mutations, indicating the sensitivity of the tool; 2) the percentage of simulation runs in which the disease gene passes GWSL, indicating the statistical power of the tool; and 3) the number of significant autosomal candidate genes, indicating the specificity of the tool. Furthermore, the performance of GRIPT was compared with four popular cohort analysis tools, including the Mendelian disease gene finder, VAAST2, and three group-wise association tests, the CMC (burden test), SKAT and KBAC (kernel model), on the same datasets.

#### The sensitivity and specificity of GRIPT under the AR and AD models

To test the performance of GRIPT in identifying AR disease gene, *RPE65* was used as an example. *RPE65* is a well-studied gene with mutations known to cause AR Leber congenital amaurosis (LCA) and Retinitis Pigmentosa (RP) [20-22]. The performance of the four tests was summarized in Figure 3 and Supplementary table S1. Figure 3A-C and table S1 demonstrate that GRIPT has great sensitivity and specificity in detecting *RPE65*, even when the proportion of *RPE65* patients was very low, mimicking the scenario of patient cohort with high locus heterogeneity. When the *RPE65* patient proportion was as low as 0.5%, GRIPT ranked *RPE65* on average sixth, achieving 66.67% power. When the *RPE65* patient proportion reached ≥ 1%, GRIPT ranked *RPE65* first in all trials with 100% power. Across the range of *RPE65* patient proportions, GRIPT identified on average three significant candidates per simulation. In contrast, with a low proportion of *RPE65* patients, the other four algorithms had significantly lower sensitivity and power than GRIPT (WRST, p-value see Supplementary table S1). For example, when the *RPE65* patient proportion was ≤ 1%, the powers of the other four tests were ≤ 10% and the mean rank of *RPE65* was between 38 and 3068. Each of the other four methods identified on average zero or one significant candidate gene.

**Figure 3.**
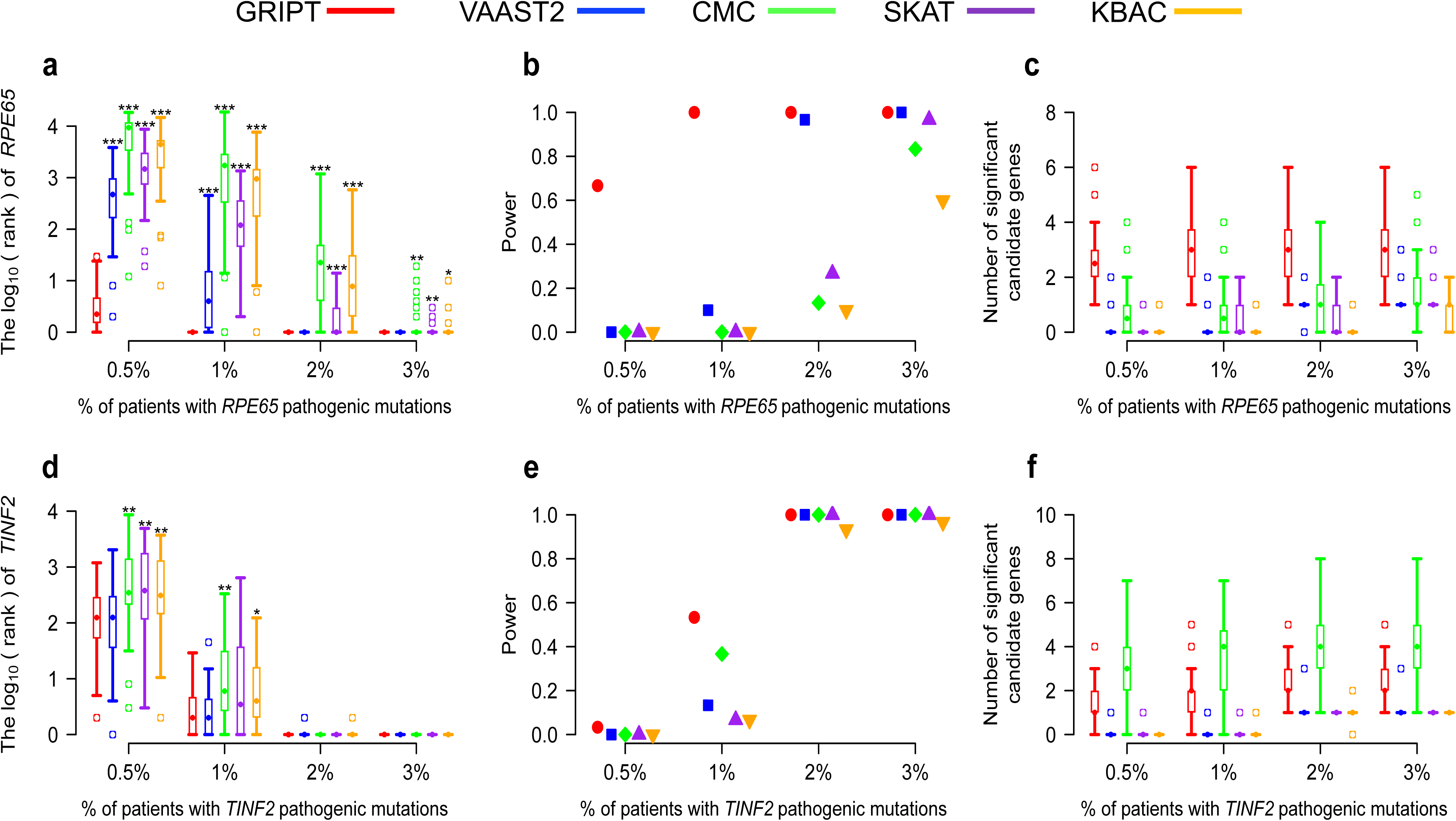
Simulation analysis of GRIPT, VAAST2, CMC, SKAT and KBAC under the AR and AD models. The AR and AD models were tested with 0.5%, 1%, 2%, and 3% of patients carrying the pathogenic mutations of *RPE65* or *TINF2,* respectively. The patient cohort size was 600. The control cohort size was 5000. The performance of GRIPT, VAAST2, CMC, SKAT and KBAC are shown in red, blue, green, purple, and orange, respectively. a) The ranking of *RPE65* under the AR model were shown in boxplot. b) The power of the five tools were measured as the proportion of simulation runs in which *PRE65* passed the GWSL shown in dot plot. c) The number of significant autosomal candidate genes under the AR model were shown in boxplot. d) The ranking of *TINF2* under the AD model. e) The power of the five tools for *TINF2*. f) The number of significant autosomal candidates under the AD model. The rankings of *RPE65/TINF2* generated by GRIPT were compared to those generated by the other four methods respectively with one-tailed WRST. The methods that generated significantly worse ranking than GRIPT were marked with ‘* ‘if p-value < 0.05, ‘** ‘if p-value < 0.01, and ‘*** ‘if p-value < 0.001.

In parallel, the performance of GRIPT in identifying AD disease gene was tested using *TINF2* as an example. *TINF2* is a known, disease-causing gene of AD Revesz syndrome and Dyskeratosis congenital [23-25]. As shown in Figure 3D-F and table S1, GRIPT lacked power when the *TINF2* patient proportion was very low, but its performance was greatly improved as the *TINF2* patient proportion increased. Specifically, as *TINF2* patient proportion increased from 0.5% to 1%, the power of GRIPT increased from 3.33% to 53.33%. When the *TINF2* patient proportion reached ≥ 2%, TINF2 was always ranked first by GRIPT with 100% power. On average, GRIPT identified about two significant candidate genes. In comparison, the other four methods had significantly worse performance than GRIPT (WRST, p-value see Supplementary table S1). For example, when *TINF2* patient proportion increased from 0.5% to 1%, the power of VAAST2 increased from 0% to 13.33%, CMC from 0% to 36.67%, SKAT from 0% to 6.67%, and KBAC from 0% to 6.67%.

#### Benchmark on 400 randomly selected known disease genes

To further expand the evaluation of GRIPT, we performed simulation using 400 Mendelian disease-causing genes randomly selected from the OMIM database, including 200 AR and 200 AD disease genes. For each gene, we simulated the patient cohorts with a size of 600 and used the same simulated control cohort with a size of 5000. The results were summarized in Figure 4 and Supplementary table S2.

**Figure 4.**
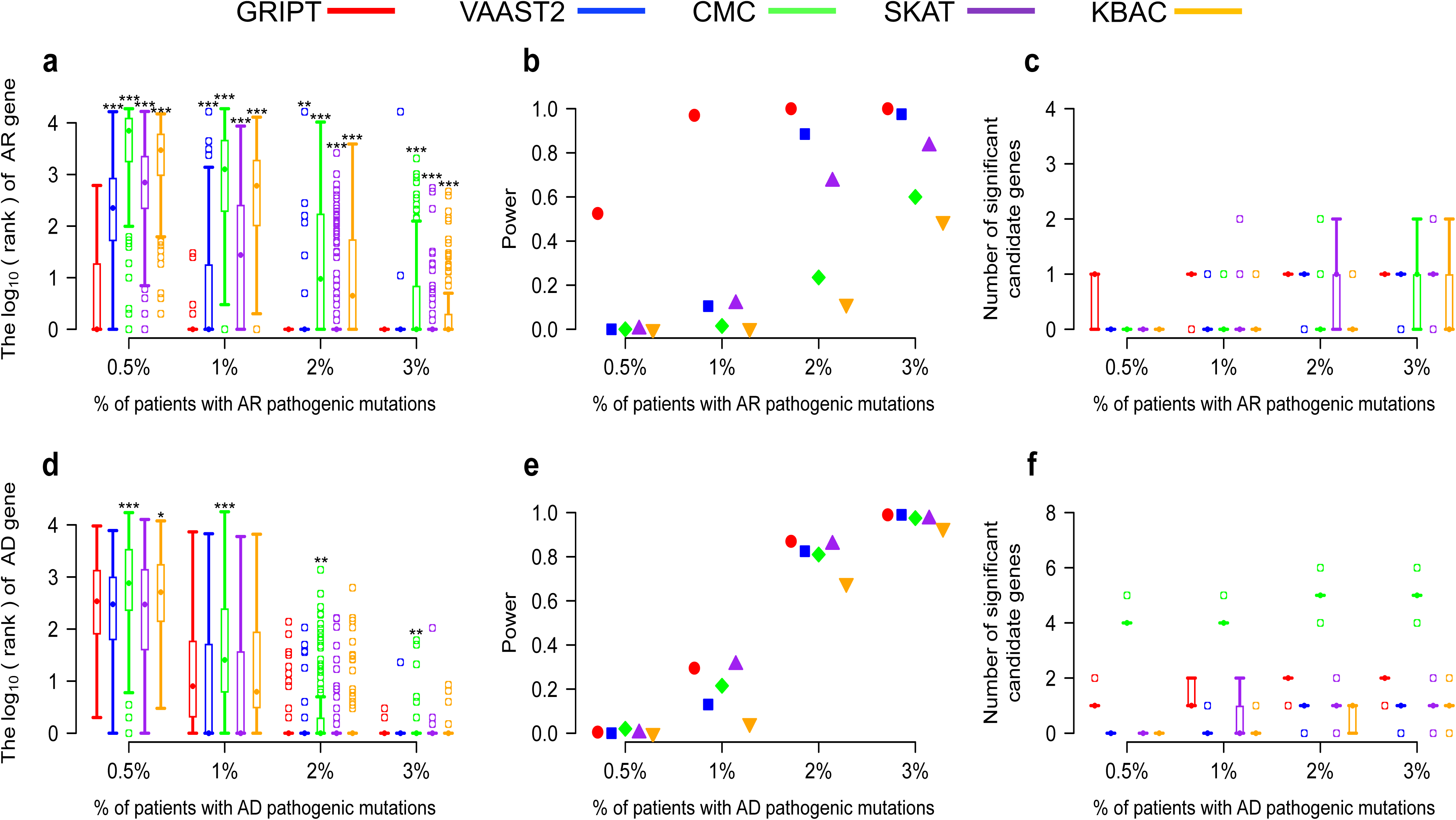
Benchmark of GRIPT, VAAST2, CMC, SKAT and KBAC on 400 Mendelian disease genes. The AR and AD models were tested with 0.5%, 1%, 2%, and 3% of patients carrying the pathogenic mutations of each of 200 AR genes and each of 200 AD genes, respectively. The patient cohort size was 600. The control cohort size was 5000. The performance of GRIPT, VAAST2, CMC, SKAT and KBAC are shown in red, blue, green, purple, and orange, respectively. a) The ranking of 200 AR genes. b) The power of the five tests for 200 AR genes. c) The number of significant autosomal candidates under the AR model. d) The ranking of 200 AD genes. e) The power of the five tests for 200 AD genes. f) The number of significant autosomal candidates under the AD model. The rankings of AR/AD genes generated by GRIPT were compared to those generated by the other four methods respectively with one-tailed WRST. The methods that generated significantly worse ranking than GRIPT were marked with ‘* ‘if p-value < 0.05, ‘** ‘if p-value < 0.01, and ‘*** ‘if p-value < 0.001.

Consistent with the results for *RPE65*, GRIPT showed outstanding sensitivity and specificity in detecting the 200 AR genes even when the proportion of patients attributed to the same disease gene was very low (Figure 4A-C). Consistently, VAAST2, CMC, SKAT and KBAC showed significantly worse performance than GRIPT when patient cohort had high locus heterogeneity (Figure 4A-C, WRST, p-value see Supplementary table S2). When the proportion of patients attributed to the same disease gene was as low as 0.5%, the disease genes were ranked on average 24^th^ by GRIPT achieving 52.5% power, whereas the other four methods had 0% power. When the patient proportion equaled to 1%, the disease genes were ranked on average first by GRIPT with 97% power. In contrast, the power of the other four methods were between 0.5% and 11.5%. When the patient proportion reached ≥ 2%, the disease genes were always ranked first by GRIPT with 100% power. In comparison, the power of the four methods were between 11.5% and 97.5%. Across the range of patient proportions, GRIPT identified on average one significant candidate gene compared to zero or one candidate by each of the other four methods.

Consistent to the results of *TINF2*, the overall performance of GRIPT was better than or comparable to the other four methods in detecting the 200 AD genes (WRST, p-value see Supplementary table S2). When the proportion of patients attributed to the same disease gene was ≤ 1%, GRIPT and the other four tests have very low power, i.e. ≤ 29.5% for GRIPT, ≤ 13% for VAAST2, ≤ 21.5% for CMC, ≤ 31% for SKAT, ≤ 4.5% for KBAC (Figure 4D-F). When the patient proportion attributed to the same gene increased to 2%, the disease genes were ranked on average third by GRIPT with 87% power. In comparison, the power of the other four tests were between 68% and 85.5%. When the patient proportion reached 3%, the disease genes were ranked first in 97.5% of simulations by GRIPT with 99% power. Comparably, the power of the other four tests increased to 93% - 99%. Across the range of patient proportions, on average one to two significant candidate genes were identified by GRIPT compared to between zero and five candidates by the other four methods.

### Simulations suggest GRIPT is highly robust

The performance of case-control cohort analysis can be potentially impacted by several confounding factors, such as patient cohort size, population stratification, and the cutoff of variant filtering frequency, and the control cohort size. To assess their impact, we performed simulations using *RPE65* and *TINF2* as examples under the AR and AD models respectively, and compared GRIPT with VAAST2, CMC, SKAT and KBAC using the same datasets under each scenario. In addition, we tested the effect of different variant score systems on the performance of GRIPT.

#### The sample size of the patient cohort

We simulated the patient cohorts in a range of sizes, i.e. 50, 100, 300, 600, and 800, with 2% of patients carrying the pathogenic mutations of the same disease genes, and control cohorts with a size of 5000. The results were summarized in Figure 5 and Supplementary table S3.

**Figure 5.**
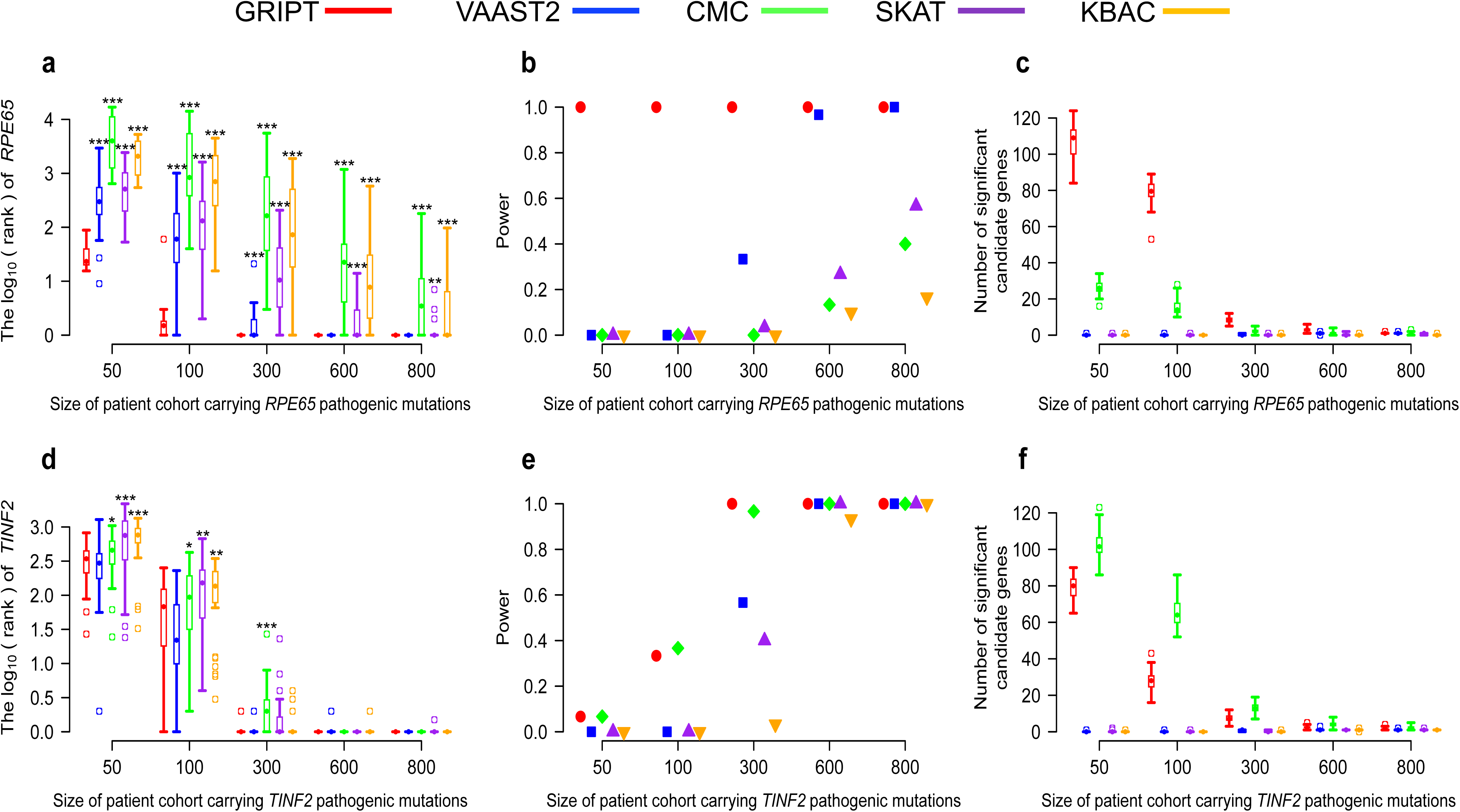
The impact of patient cohort sizes. The patient cohort sizes were tested at 50, 100, 300, 600 and 800. The control cohort size was set at 5000. The percentage of patients carrying the pathogenic mutations of *RPE65* or *TINF2* was set at 2%. The performance of GRIPT, VAAST2, CMC, SKAT and KBAC are shown in red, blue, green, purple, and orange, respectively. a) The ranking of *RPE65* under the AR model. b) The power of the five tests for *RPE65*. c) The number of significant autosomal candidates under the AR model. d) The ranking of *TINF2* under the AD model. e) The power of the five tests for *TINF2*. f) The number of significant autosomal candidates under the AD model. The rankings of *RPE65/TINF2* generated by GRIPT were compared to those generated by the other four methods respectively with one-tailed WRST. The methods that generated significantly worse ranking than GRIPT were marked with ‘* ‘if p-value < 0.05, ‘** ‘if p-value < 0.01, and ‘*** ‘if p-value < 0.001.

As shown in Figure 5A-C, under the AR model, GRIPT maintains high sensitivity for patient cohorts with a variety of sizes and high locus heterogeneity although its specificity decreased for small patient cohorts with high locus heterogeneity. In comparison, the other four methods performed significantly worse than GRIPT under the same situations (WRST, p-value see Supplementary table S3). Specifically, as the patient cohort size increased from 50 to 300 with 2% of patients carrying the *RPE65* pathogenic mutations, the mean rank of *RPE65* increases from 31 to 1 by GRIPT with 100% power. The number of significant candidates identified by GRIPT decreased from 107 to 8. When the patient cohort size reached ≥ 300, GRIPT always ranked *RPE65* first with 100% power. The average number of significant candidates decreased to between one and eight. In contrast, the power of the other four methods was 0% when the patient cohort size < 300. When the patient cohort size reached ≥ 300, the power was 33.33%-100% for VAAST2, 0%-40% for CMC, 3%-56.67% for SKAT, and 0%-16.67% for KBAC. And the average number of significant candidates identified by each of the four methods was between 0 and 26.

Under the AD model, when patient cohort was small and had high locus heterogeneity, GRIPT had low sensitivity and specificity, but its performance was greatly improved as the patient cohort size increased (Figure 5D-F). The other four methods performed comparably or significantly worse under the same scenarios (Figure 5D-F, WRST, p-value see Supplementary table S3). Specifically, when the patient cohort size increased from 50 to 100 with 2% of patients attributed to *TINF2*, the power of GRIPT increased from 6.67% to 33.33% and the average number of significant candidates decreased from 79 to 28. When the patient cohort size increased to ≥ 300, *TINF2* was ranked on average first by GRIPT with 100% power. The average number of significant candidates by GRIPT was between two and eight. In comparison, when the patient cohort size < 300, the power increased from 6.67% to 36.67% for CMC and remained at 0% for VAAST2, SKAT and KBAC. When the patient cohort size reached ≥ 300, the power was between 3.33% and 100% for the four tests. The average number of significant candidates by each of the four tests was between 0 and 103.

#### Population stratification of cohorts

It was observed that the variant spectrum of a disease-gene is different among populations with different ethnicities and that high population stratification could impair the performance of cohort analysis [16]. To test the impact of population stratification on GRIPT, we simulated patient cohorts as an admixture of African and Latino individuals and control cohorts with Latino individuals only, based on the allele frequency in ExAC database with corresponding ethnicity (see Methods). The unmatched proportion between case and control cohorts were simulated at 0%, 20%, 40%, 60%, 80% and 100%. The size of patient cohort was set at 500 and the control cohort at 5000. The proportion of patients carrying the pathogenic mutations of the same gene was set at 1%. The results were summarized in Figure 6 and Supplementary table S4.

**Figure 6.**
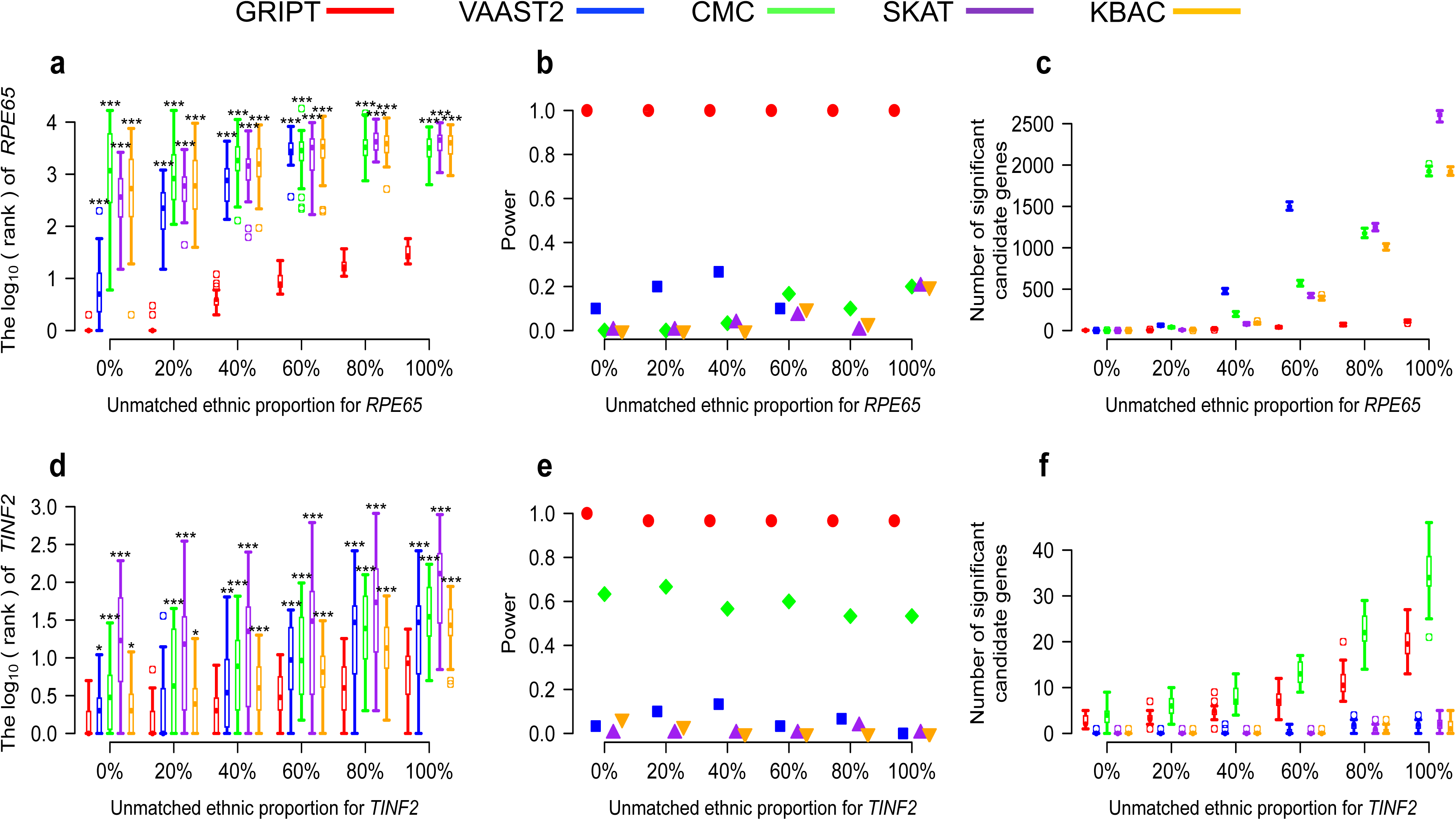
The impact of population stratification. The unmatched proportions between patient cohort and control cohort were tested at 0%, 20%, 40%, 60%, 80% and 100%. The percentage of patients carrying the *RPE65* or *TINF2* pathogenic mutations was set at 1%. The patient cohort size was 500. The control cohort size was 5000. The performance of GRIPT, VAAST2, CMC, SKAT and KBAC are shown in red, blue, green, purple, and orange, respectively. a) The ranking of *RPE65* under the AR model. b) The power of the five tests for *RPE65*. c) The number of significant autosomal candidates under the AR model. d) The ranking of *TINF2* under the AD model. e) The power of the five tests for *TINF2*. f) The number of significant autosomal candidates under the AD model. The rankings of *RPE65/TINF2* genes generated by GRIPT were compared to those generated by the other four methods respectively with one-tailed WRST. The methods that generated significantly worse ranking than GRIPT were marked with ‘* ‘if p-value < 0.05, ‘** ‘if p-value < 0.01, and ‘*** ‘if p-value < 0.001.

As shown in Figure 6A-F, the sensitivity and specificity of GRIPT slightly decreased as unmatched ethnicity proportion between cases and controls increased. However, GRIPT is significantly less affected by population stratification than the other four methods even when patient cohort had high locus heterogeneity (WRST, p-value see Supplementary table S4). Specifically, under the AR model, as the unmatched ethnicity proportion between patients and controls increased from 0% to 100% (namely, from the completely matched to the completely unmatched), the mean rank of *RPE65* dropped from 1 to 32 by GRIPT but always with 100% power (Figure 6A-C). Specificity was reduced as the average number of significant candidate genes increased from 2 to 111 (Figure 6A-C). In comparison, the powers of CMC, SKAT and KBAC were between 0% and 20%. The average number of significant candidate genes increased from 1 to 1929 for CMC, from 0 to 2603 for SKAT, and from 0 to 1921 for KBAC. In addition, as the unmatched ethnicity proportion increased, the running time for VAAST2 dramatically increased (e.g. needs 120-240 hours with 5 parallel CPUs to finish one simulation run), VAAST2 was only tested for the unmatched ethnicity proportion ranging from 0% to 60%. Under those scenarios, the power of VAAST2 was between 10% and 26.7%. The average number of significant candidate genes identified by VAAST2 increased from 0 to 1502.

Under the AD model, GRIPT is also significantly less affected by population stratification (WRST, p-value see Supplementary table S4). As the unmatched ethnicity proportion increased from 0% to 100%, the mean rank of *TINF2* dropped from two to nine by GRIPT with 96.67%-100% power (Figure 6D-F). The mean number of significant candidate genes increased from 3 to 19. In comparison, the mean rank of *TINF2* dropped from 3 to 75 for VAAST2, from 7 to 57 for CMC, and from 44 to 166 for SKAT, and from 3 to 33 for KBAC. The power was 0%-13.33% for VAAST2, 53.33%-66.67% for CMC, 0%-3.33% for SKAT, and 0%-6.67% for KBAC. The average number of significant candidate genes increased from zero to five for VAAST2, from 4 to 35 for CMC, from zero to two for SKAT, and from zero to one for KBAC. (Figure 6D-F).

#### Variant frequency filtering

Mendelian disease-causing mutations are expected to be very rare in the population, and common human variants are likely benign for rare Mendelian diseases. Therefore, to reduce the analysis/computation complexity, variants from WES are conventionally first filtered out common human genome variants based on allele frequency in large database of human genome variants, e.g. gnomAD and ExAC. To mimic this scenario, the above patient and control cohorts were simulated using the variants whose maximum population frequency ≤ 0.5% in ExAC database for the AR model, and whose maximum population frequency ≤ 0.01% for the AD model. Here, we examined the impact of a relaxed (i.e. higher) frequency filtering cutoff on the disease gene identification methods. We simulated the WES data of patient and control cohorts using a range of variant frequency cutoffs respectively: ≤ 0.5%, ≤ 1% and ≤ 2% for the AR model, and ≤ 0.01%, ≤ 0.5% and ≤ 1% for the AD model. The proportion of patients attributed to the same gene was set at 1%. The size of patient cohort was set at 600 and control cohort at 5000. The results show that inclusion of more variants/noise per individual by using higher frequency filtering cutoff had little impact on GRIPT’s performance under the AR model, but it reduced its power under the AD model. The performance of the other four methods were largely compromised and were significantly worse than or comparable to that of GRIPT (Figure 7A-F, Supplementary table S5).

**Figure 7.**
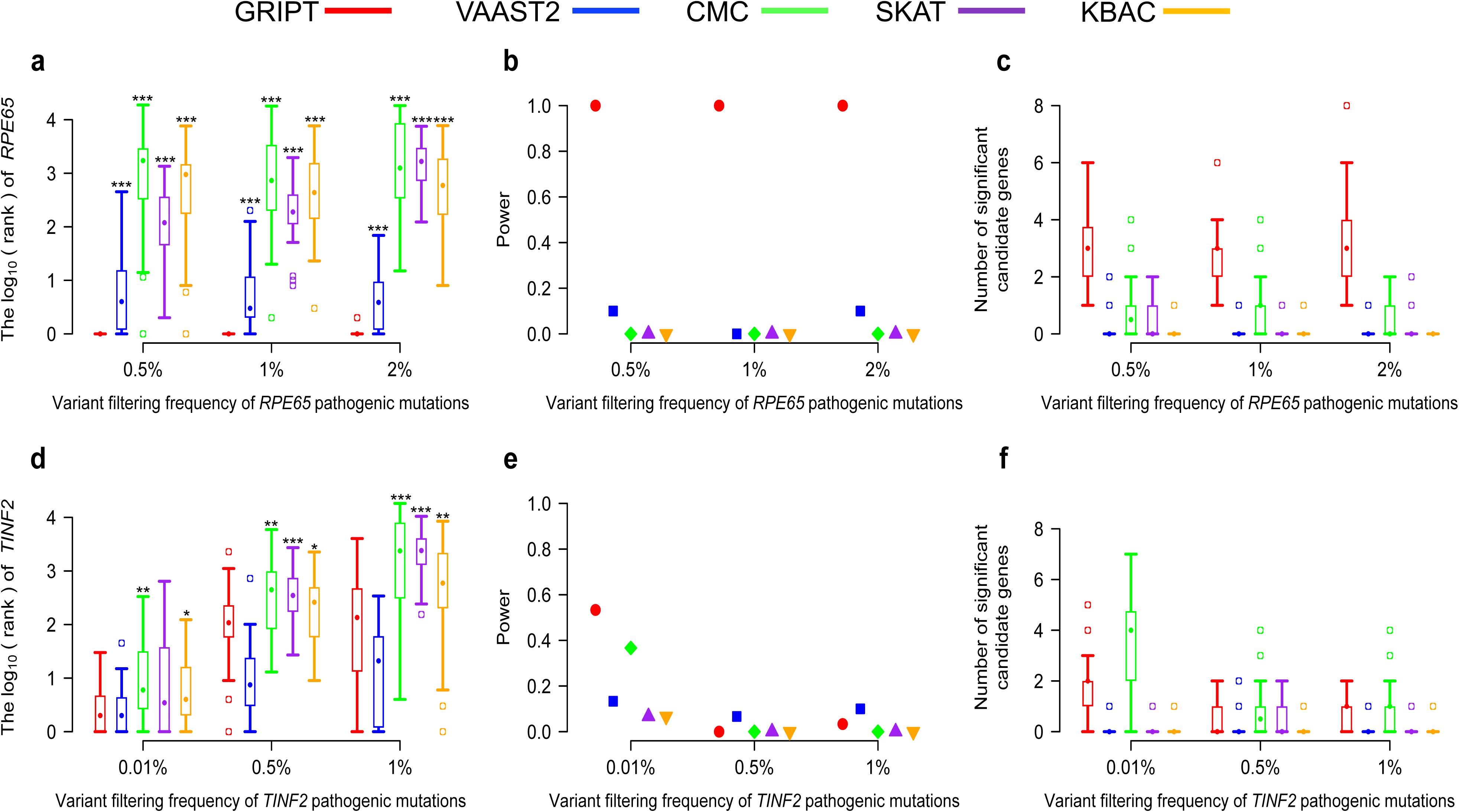
The impact of variant frequency filtering. The cutoff of variant filtering frequency was tested at 0.5%, 1%, and 2% under the AR model, and at 0.01%, 0.5%, and 1% under the AD model. The percentage of patients carrying the *RPE65* or *TINF2* pathogenic mutations was set at 1%. The patient cohort size was 600. The control cohort size was 5000. The performance of GRIPT, VAAST2, CMC, SKAT and KBAC are shown in red, blue, green, purple, and orange, respectively. a) The ranking of *RPE65* under the AR model. b) The power of the five tests for *RPE65*. c) The number of significant autosomal candidates under the AR model. d) The ranking of *TINF2* under the AD model. e) The power of the five tests for *TINF2*. f) The number of significant autosomal candidates under the AD model. The rankings of *RPE65/TINF2* generated by GRIPT were compared to those generated by the other four methods respectively with one-tailed WRST. The methods that generated significantly worse ranking than GRIPT were marked with ‘* ‘if p-value < 0.05, ‘** ‘if p-value < 0.01, and ‘*** ‘if p-value < 0.001.

Specifically, under the AR model, as the frequency filtering cutoff increased from 0.5% to 2%, GRIPT ranked *RPE65* first in 98.89% of the simulations, always achieving 100% power. The mean number of significant candidate genes was about three (Figure 7A-C). In contrast, the ranking of *RPE65* by the other four tests was largely decreased, with ≤ 10% power for VAAST2, 0% power for CMC, SKAT and KBAC. Under the AD model, as the variant frequency cutoff increased from 0.01% to 1%, the average rank of *TINF2* dropped from 5 to 590 by GRIPT with power decreasing from 53.33% to around 3%. The average number of significant candidate genes was between zero to two (Figure 7D-F). The power of VAAST2 decreased from 13.33% to 10%, CMC from 36.67% to 0%, SKAT from 6.67% to 0% for SKAT, and KBAC from 6.67% to 0%.

#### The effect of the control cohort size

Theoretically, the variant spectrum of a gene in a large control cohort should be less biased and closer to the true distribution than that in a small control cohort. Thus, large control cohorts can better serve as the control/baseline, for example, to exclude the genes with large numbers of rare benign variants in population. To test the effect of control cohort size, we simulated smaller control cohorts with a size of 600 and used the previous case cohorts with a size of 600 to repeat the analysis. The results were summarized in Figure 8 and Supplementary table S6.

**Figure 8.**
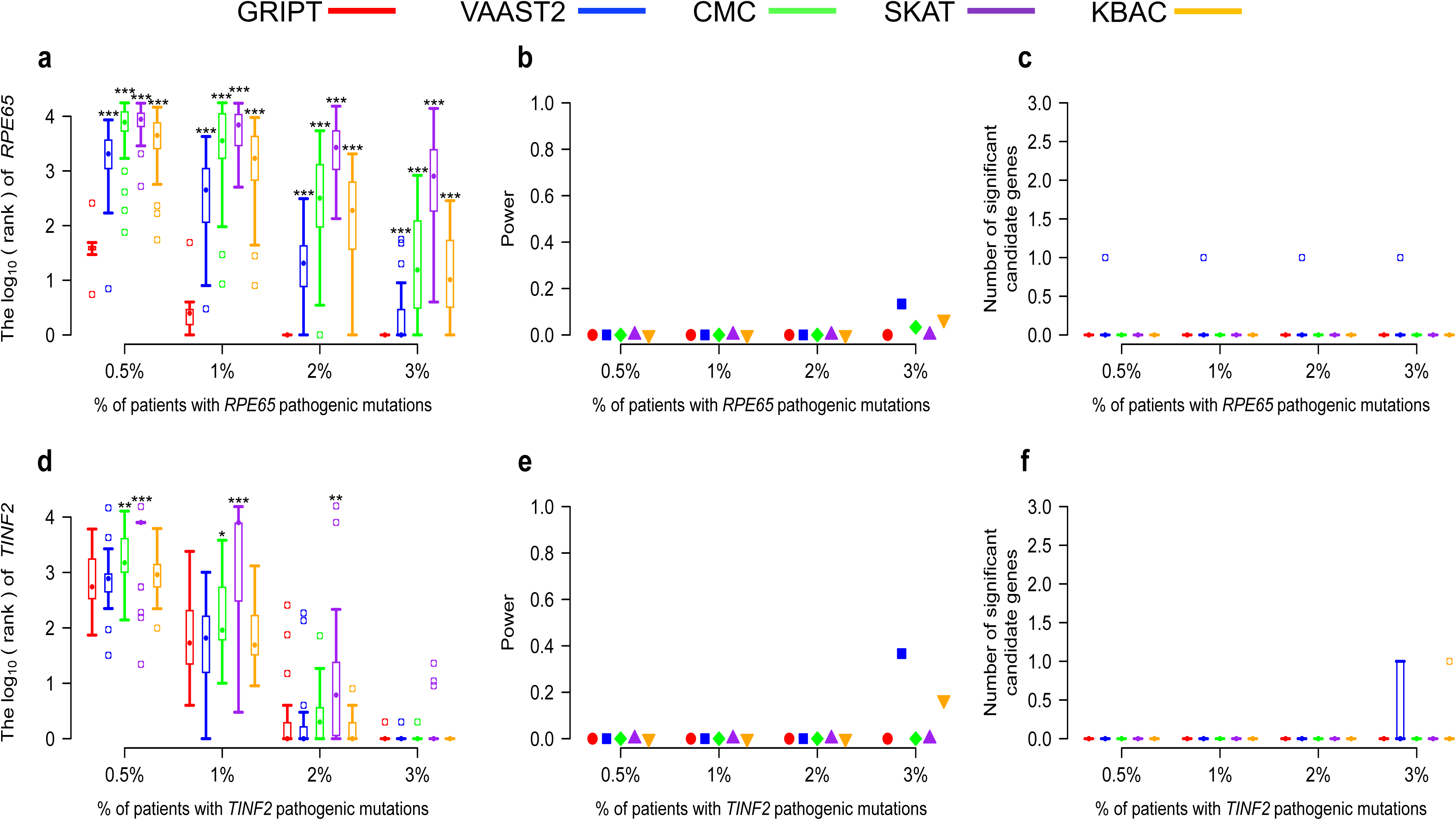
The effect of control cohort sizes. The AR and AD models were tested with 0.5%, 1%, 2%, and 3% of patients carrying the pathogenic mutations of *RPE65* or *TINF2*, respectively. The patient cohort size was 600. The control cohort size was 600. The performance of GRIPT, VAAST2, CMC, SKAT and KBAC are shown in red, blue, green, purple, and orange, respectively. a) The ranking of *RPE65* under the AR model. b) The power of the five tools for *RPE65*. c) The number of significant autosomal candidates under the AR model. d) The ranking of *TINF2* under the AD model. e) The power of the five tools for *TINF2*. f) The number of significant autosomal candidates under the AD model. The rankings of *RPE65/TINF2* generated by GRIPT were compared to those generated by the other four methods respectively with one-tailed WRST. The methods that generated significantly worse ranking than GRIPT were marked with ‘* ‘if p-value < 0.05, ‘** ‘if p-value < 0.01, and ‘*** ‘if p-value < 0.001.

Under the AR model, GRIPT remained sensitive in ranking *RPE65*. When the *RPE65* patient proportion increased from 0.5% to ≥ 2%, the mean rank of RPE65 increased from 45 to 1. However, the p-value of *RPE65* did not pass the GWSL in any of the simulations, showing GRIPT with 0% power. Consistent to the results with larger control cohort, the other four tools performed significantly worse than GRIPT (Figure 8A-C, WRST, p-value see Supplementary table S6). For example, when the *RPE65* patient proportion equaled to 1%, the mean rank of RPE65 was 981 for VAAST2, 6243 for CMC, 7611 for SKAT and 2892 for KBAC. Similarly, the p-values of *RPE65* from the other four tests did not pass the GWSL for the majority of the simulations either, shown as the test power below 13.33%.

Under the AD model with the small control cohorts, the rankings of *TINF2* by GRIPT and the other four methods were consistent to that with the large control cohorts (Figure 8D-F, Supplementary table S6). The five methods gave *TINF2* a low ranking when the *TINF2* patient proportion was low. But the ranking of *TINF2* rose as the *TINF2* patient proportion increased. When the TINF2 patient proportion increased to 3%, all five methods ranked *TINF2* to the top. However, similar to the results under the AR model, the p-value of *TINF2* by the five methods did not pass the GWSL in the majority of the simulations under the AD model, shown as the power below 36.67% (Figure 8D-F).

#### The effect of different variant scoring systems

To test whether the performance of GRIPT will be affected by different variant score systems, besides CADD score, we applied the DANN and REVEL scores to annotate the variant scores in GRIPT respectively, and repeated the aforementioned analyses. DANN scoring system shares the same feature set and training data as CADD (which was trained with a linear kernel support vector machine, SVM) but was trained with a non-linear deep neural network. DANN achieves about a 19% relative reduction in the error rate and about a 14% relative increase in the area under the curve (AUC) metric over CADD’s SVM methodology [26]. REVEL is an ensemble method for predicting the pathogenicity of missense variants by integrating the individual tools, including MutPred, FATHMM, VEST, PolyPhen, SIFT, PROVEAN, MutationAssessor, MutationTaster, LRT, GERP, SiPhy, phyloP, and phastCons. REVEL outperformed (p < 10^-12^) individual tools and seven ensemble methods (i.e. MetaSVM, MetaLR, KGGSeq, Condel, CADD, DANN, and Eigen) in analyzing independent test sets, and also showed the best performance for distinguishing pathogenic from rare neutral variants with allele frequencies <0.5% [27]. As shown in the Supplementary Figure S2-S5 and Supplementary Table S2-S5, the benchmark analysis with 400 AR and AD genes, the analyses of the impacts of patient cohort size, population stratification, and variant frequency filtering all showed that the results based on DANN and REVEL scores are consistent with the previous results based on CADD score. The consistency based on different variant score systems demonstrated the reliability and robustness of the statistic test framework of GRIPT.

#### Comparison to the traditional GWAS single variant test

To compare the performance of GRIPT with the traditional GWAS single variant test, we simulated the basic scenario with 0.5%-3% of patients carrying the pathogenic mutations of *RPE65* and *TINF2* respectively, and applied GRIPT and Fisher’s exact test to the data. As shown in Figure 9 and Supplementary table S1, Fisher’s exact test performed much worse than GRIPT. Under the AR model, when the *RPE65* patient proportion was 0.5%, *RPE65* was ranked on average sixth by GRIPT with 66.67% power. When the *RPE65* patient proportion was ≥ 1%, *RPE65* was always ranked first by GRIPT with 100% power. In contrast, the average ranking of *RPE65* by Fisher’s exact test was in the range of 890 to 32000, always with 0% power. Under the AD model, as *TINF2* patient proportion increased from 0.5% to 1%, the power of GRIPT increased from 3.33% to 53.33%. When the *TINF2* patient proportion was ≥ 2%, GRIPT always ranked TINF2 first with 100% power. In comparison, as the proportion of *TINF2* patients increased, the average ranking of *TINF2* by Fisher’s exact test was improved from 12675^th^ to 23^th^, but the test power remained at 0%. The reasons may be: 1) GRIPT is a gene-wise test that ranks the functional effects of variants and incorporates the Mendelian inheritance models to compute the gene score. In contrast, the traditional single variant test considers one variant in a gene each time, and is mainly based on the allele frequency difference between cases and controls. Thus, the single variant test does not have sufficient power to detect the heterogeneous rare deleterious variants in Mendelian disease cohorts, although it might be suitable for common complex diseases. 2) the multiple test correction requests a much more stringent p-value cutoff for the single variant test than the gene-wise GRIPT due to the larger number of tests applied in the single variant test than in GRIPT (i.e. variants vs. genes).

**Figure 9.**
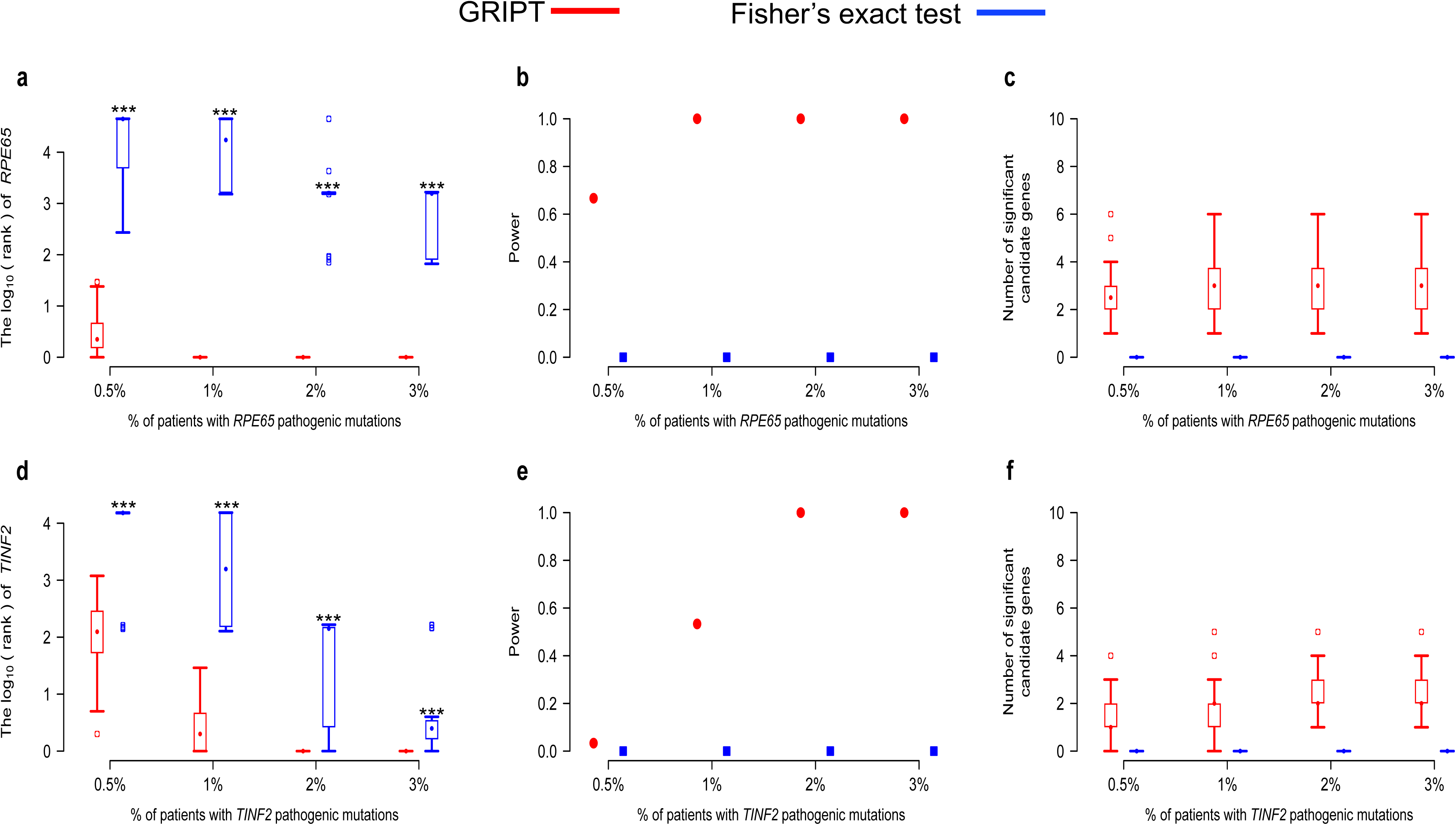
The comparison of the performance of Fisher’s exact test with GRIPT. The AR and AD models were tested with 0.5%, 1%, 2%, and 3% of patients carrying the pathogenic mutations of *RPE65* or *TINF2,* respectively. The patient cohort size was 600. The control cohort size was 5000. The performance of GRIPT and Fisher’s exact test are shown in red and blue, respectively. a) The ranking of *RPE65* under the AR model. b) The power of the two tests for *RPE65*. c) The number of significant autosomal candidate genes under the AR model. d) The ranking of *TINF2* under the AD model. e) The power of the two tests for *TINF2*. f) The number of significant autosomal candidates under the AD model. The rankings of *RPE65/TINF2* generated by GRIPT were compared to those generated by the other four methods respectively with one-tailed WRST. The methods that generated significantly worse ranking than GRIPT were marked with ‘* ‘if p-value < 0.05, ‘** ‘if p-value < 0.01, and ‘*** ‘if p-value < 0.001.

### Analysis of real patient cohort data display GRIPT’s excellent performance

To further validate the performance of GRIPT, we applied it to real WES data of three different patient cohorts respectively, including a Leber’s congenital amaurosis (LCA) cohort, a Retinitis pigmentosa (RP) cohort, and a congenital disorder of glycosylation (CDG) cohort. Both the LCA cohort and RP cohort were composed of the patients carrying the pathogenic mutations of different genes, and the proportion of patients attributed to each disease gene was small. Furthermore, the patient ethnicity of the LCA cohort or RP cohort was an admixture of Caucasian, African American, Latino, and Asian. Whereas, the CDG cohort was composed of the patients all attributed to *PGM3* from two families. The performance of GRIPT was also compared with VAAST2, CMC, SKAT and KBAC on the same datasets.

#### The LCA cohort

LCA is a genetic heterogeneous disease and can be caused by mutations in at least 22 genes (http://www.sph.uth.tmc.edu/RetNet, accessed as September 3rd, 2017). We performed WES on 115 sporadic LCA patients. As LCA is a rare Mendelian disorder, variants with maximum population allele frequency > 0.5% were filtered out based on the allele frequency in the large public databases of normal populations (i.e. 1000 genome, dbSNP, ESP6500, ExAC, gnomAD) and an internal database. We only focused on rare protein-changing variants including nonsense variants, splicing donor/acceptor variants, missense variants, and small INDELs, since they are more likely to be the disease-causing mutations. One previously simulated control cohort (n=5000) was used as the control cohort for these tests.

GRIPT showed high sensitivity for the LCA cohort with high locus and ethnicity heterogeneity. It successfully detected the disease gene that only accounted for ≤ 1% of the patients. Specifically, the first nine candidate genes ranked by GRIPT were all known retinal disease genes (Table 1). Among a total of 203 significant candidates, 19 genes were known disease genes, each of which accounted for 0.87%-6.09% (one to seven patients) of the cohort. Most interestingly, GRIPT was able to identify novel retinal disease genes, i.e. *POMGNT1* (p = 2.81 × 10^-10^) and *MFSD8* (p = 2.81 × 10^-10^). *POMGNT1* was a gene causing non-syndromic RP newly discovered in 2016 [28], and accounted for one patient of this cohort, who carried a stop-gain mutation and a missense mutation in *POMGNT1*. Mutations in *MFSD8* have been linked to Macular Dystrophy recently [29] and accounted for one patient of the LCA cohort, who carried a splice donor mutation and a missense mutation in *MFSD8*.

**Table 1.**
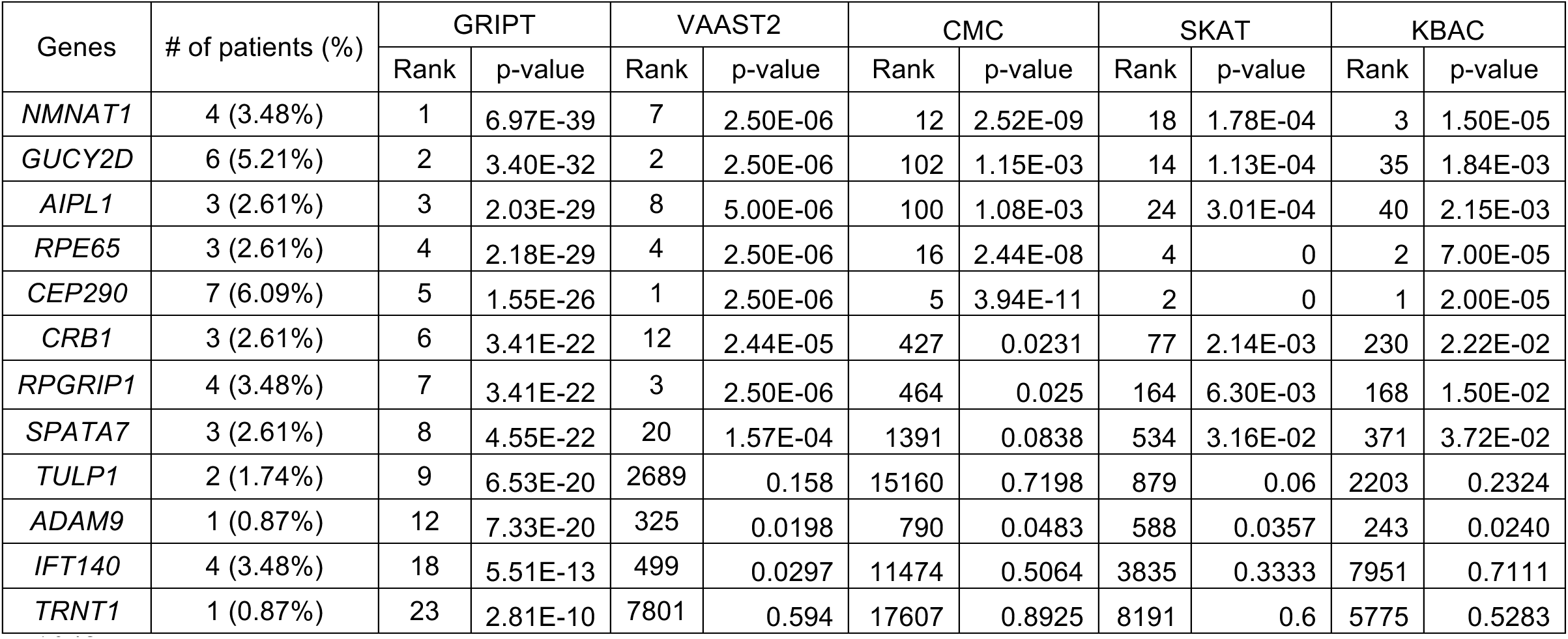
Known disease genes were given high ranks and significant P-values by GRIPT in a LCA cohort. The listed genes are the correctly identified retinal disease genes among the top 20 candidate genes by GRIPT in the LCA cohort. Parameters: 115 cases, 5000 controls, the AR inheritance model.

In comparison, the other tools lacked power in detecting the disease genes accounting for small proportions of this cohort. A total of 7 significant candidates were identified by VAAST2, 27 by CMC, 6 by SKAT, and 1 by KBAC. Among them, 5 genes by VAAST2 were known disease genes, 3 genes by CMC, 2 genes by SKAT, and 1 genes by KBAC, each of which accounted for 2.61%-6.09% (three to seven patients) of the cohort. However, none of these known genes were the recently identified novel retinal disease genes.

#### *The RP coho*rt

RP is an inherited retinal disease with even greater genetic heterogeneity compared to LCA. So far, mutations in more than 65 genes were found to cause the disease (http://www.sph.uth.tmc.edu/RetNet, accessed by September 3rd, 2017). WES was performed for 154 sporadic RP patients. After filtering, the WES data of the real patient cohort and a simulated control cohort (n=5000) were subjected to analysis. GRIPT again showed excellent power in identifying low frequency disease genes underlying the cohort with high locus and ethnicity heterogeneity. As shown in Table 2, eight genes whose rankings ranged from first to eleventh by GRIPT were known retinal disease genes. Among the 157 significant candidates (p < 2.7e-6) identified by GRIPT, 17 are known disease genes, each of which explained 0.649%-8.44% (1 to 13 patients) of the cohort. Furthermore, GRIPT was able to identify three novel retinal disease genes recently published, i.e. *POMGNT1* (p = 3.95 × 10^-15^)*, TRNT1* (p = 6.25 × 10^-8^) and *HGSNAT* (p=2.10 × 10^-7^). Mutations in *POMGNT1* [28] accounted for two patients of the cohort, who carried two different homozygous missense mutations. Mutations in *HGSNAT*, a gene causing nonsyndromic RP[30], explained two patients in this cohort. One patient carried two missense mutations, and the other carried a disruptive inframe deletion and a missense mutation. Mutations in *TRNT1*, a gene causing RP and erythrocytic microcytosis[31], accounted for one patient in the cohort, who carried a frameshift mutation and a missense mutation in *TRNT1*.

**Table 2.** Known disease genes were given high ranks and significant P-values by GRIPT in a RP cohort. The listed genes are the correctly identified retinal disease genes among the top 20 candidate genes by GRIPT in the RP cohort. Parameters: 154 cases, 5000 controls, the AR inheritance model.

| Genes | # of patients (%) | GRIPT |  | VAAST2 |  | CMC |  | SKAT |  | KBAC |  |
| --- | --- | --- | --- | --- | --- | --- | --- | --- | --- | --- | --- |
|  |  | Rank | p-value | Rank | p-value | Rank | p-value | Rank | p-value | Rank | p-value |
| *TULP1* | 3 (1.95%) | 1 | 2.87E-22 | 15 | 1.57E-04 | 878 | 0.0429 | 279 | 0.0158 | 186 | 0.0190 |
| *EYS* | 8 (5.19%) | 2 | 5.67E-18 | 2 | 2.50E-06 | 2718 | 0.1231 | 255 | 0.0143 | 504 | 0.0556 |
| *POMGNT1* | 2 (1.30%) | 5 | 3.95E-15 | 854 | 0.0396 | 12243 | 0.5391 | 8407 | 0.6 | 7477 | 0.6315 |
| *CNGA1* | 2 (1.30%) | 6 | 3.95E-15 | 73 | 2.85E-03 | 18620 | 0.9801 | 4650 | 0.375 | 4150 | 0.3769 |
| *RDH5* | 2 (1.30%) | 7 | 3.95E-15 | 2430 | 0.119 | 2009 | 0.0924 | 1089 | 0.0769 | 900 | 0.0946 |
| *USH2A* | 13 (8.44%) | 9 | 2.27E-14 | 1 | 2.50E-06 | 44 | 6.31E-05 | 6 | 0 | 6 | 0.0001 |
| *CRB1* | 3 (1.95%) | 10 | 3.65E-11 | 114 | 4.66E-03 | 222 | 0.0063 | 428 | 0.028 | 92 | 0.0082 |
| *MERTK* | 3 (1.95%) | 11 | 6.20E-11 | 19 | 0.0003 | 6523 | 0.2699 | 76 | 0.0023 | 1325 | 0.1384 |
| *BBS4* | 2 (1.30%) | 13 | 8.51E-10 | 645 | 0.0297 | 13022 | 0.5874 | 1309 | 0.0968 | 2036 | 0.1984 |
| *MAK* | 1 (0.649%) | 17 | 6.25E-08 | 1694 | 0.0792 | 13033 | 0.5874 | 11162 | 0.75 | 3899 | 0.3573 |

In comparison, the other tools had weak power in detecting the low frequency disease genes underlying this cohort. A total of 4 significant candidate genes were identified by VAAST2, 25 by CMC, 6 by SKAT, and 2 by KBAC. Among them, 2 genes by VAAST2 were known disease genes, 0 by CMC, 1 by SKAT and 0 by KBAC, each of which accounted for 5.19%-8.44% (8 to 13 patients) of the cohort. And none of these known genes were the novel retinal disease genes recently identified.

#### The CDG cohort

The CDG cohort was composed of six patients from two families who all carry the pathogenic mutations of *PGM3* gene. The WES data were downloaded from dbGaP (phs000809.v1.p1). Thus, this cohort serves as a real data example of a genetic homogeneous disease with extremely small case cohort size from an independent external source. After filtering and annotation, the real WES data and a simulated control cohort (n = 5000) were analyzed by the five tools. GRIPT showed the highest accuracy and efficiency in analyzing this homogeneous external cohort. GRIPT correctly ranked PGM3 first (p =0), taking less than 30 minutes with one CPU. VAAST2 also ranked PGM3 first (p= 2.50 × 10^-6^) but took about 6 hours with 5 parallel CPUs. CMC ranked PGM3 11^th^ (p = 3.79 × 10^-64^) and took 2.5 hours with one CPU. The p-value of PGM3 by SKAT equals to 0 but is the same as the other 162 genes (p = 0), taking 9.3 hour with one CPU. The p-value of PGM3 by KBAC equals to 2 × 10^-6^ but is the same as the other 62 genes (p = 2 × 10^-6^), taking 7.8 hour and one CPU.

### Discussion

In this study, we developed a novel computational method named GRIPT for Mendelian disease gene discovery through analyzing the NGS data of patient-control cohorts. The null hypothesis of GRIPT is that a non-disease gene should have similar deleterious mutations load in cases and in controls. GRIPT scores and compares the deleterious mutations load of each gene in the genome between patients and controls using a composite Fisher’s test, and prioritizes the genes that have significant higher deleterious mutation loads in cases than in controls as the candidate disease genes.

Both simulation and real data tests indicate that GRIPT has great sensitivity and specificity and is highly reliable in discovering Mendelian disease genes. For example, as shown in the benchmark of 400 known disease genes, under the AR model, GRIPT ranked the disease gene first in 97.5% of the simulations for a patient cohort with a size of 600 and with only 1% of patients carrying the pathogenic mutations of the same gene. In addition, the disease gene was usually the only significant candidate gene identified by GRIPT (Figure 4A-C). Under the AD model, GRIPT ranked the disease genes in the top three in 93.5% of the simulations when 2% of patients (cohort size =600) were attributed to the same gene (Figure 3D-F). The average number of significant candidates was about two. Furthermore, the results from analysis of real patient data were consistent with the benchmark results. For the LCA cohort (size n = 115), GRIPT was able to systematically and accurately identify 19 disease genes (5 genes by VAAST2, 3 genes by CMC, 2 genes by SKAT, and 1 by KBAC). The candidates ranked from first to ninth were all real disease genes. For the RP cohort (size n = 154), GRIPT was able to accurately identify 17 genes (2 genes by VAAST2, 0 genes by CMC, 1 genes by SKAT, and 0 by KBAC) with seven of the top 10 candidates being real disease genes. Each of the disease genes identified by GRIPT only accounted for 0.649%-8.44% (1 to 13 patients) of patients in the LCA or RP cohort. Moreover, as shown in the simulation, GRIPT reached around 100% power and always ranked the genes to the top for large patient cohorts (e.g. size n ≥ 300) and/or more homogeneous patients (e.g. the same gene explaining ≥ 3% of the patients), which was also demonstrated by the analysis of the CGD cohort with a size of six and all attributed to the gene *PGM3*. Most interestingly, GRIPT was able to discover four newly reported disease genes in the analysis of real patient data. Each of these newly discovered genes only accounted for one or two (0.649%-1.3%) patients in the patient cohort. Overall, GRIPT shows the great power in discovering known and novel Mendelian disease genes. It is especially well suited to analyze diseases with high locus (and ethnicity) heterogeneity, which is a major challenge for solving the underlying genetics mechanisms of Mendelian disorders.

GRIPT is also more robust and significantly less affected by potential confounding factors than other disease gene finders. For example, GRIPT remained powerful for small patient cohorts with high locus heterogeneity. In simulation, under the AR model, for a patient cohort with a size of 100 and only two (2%) patients carrying the pathogenic mutations of the same gene, the disease gene was ranked on average third by GRIPT with 100% power. In contrast, the mean ranking of the disease gene by other tools was between ~150 and ~3300 and all with 0% power. This result was also consistent with results from real data as previously discussed. Furthermore, using higher allele frequencies as the variant filtering cutoff, which presumably adds more noise to the analysis, had little impact on the performance of GRIPT under the AR model. In the simulation, for a patient cohort with a size of 600 and with six (1%) patients attributed to the same gene, as the cutoff of variant frequency filtering increased from 0.5% to 2%, the disease gene was ranked first in 98.89% of simulations by GRIPT with 100% power. In comparison, the mean rank of the gene was between 11 and 38 by VAAST2, between 2953 and 4420 by CMC, between 269 and 2095 by SKAT, and between 1306 and 1655 by KBAC, all of which had power below 10%. More importantly, GRIPT is significantly less affected by the combined effect of population stratification and high locus heterogeneity, which occur frequently in real data and severely impair the performance of other tools as shown in the simulation and real data analysis. In the simulation of the worst-case scenario where the ethnicity of the patient cohort was completely unmatched by that of the control cohort and with only 1% of the patient cohort (with a cohort size of 500) attributed to the same disease gene under the AR model, GRIPT ranked the disease gene, on average, 32^th^ with 100% power although it generated around 107 significant candidates. In contrast, the mean ranking of the disease gene by other tools was greater than 3500 (power ≤ 20%), each of which generated more than 1500 significant candidates. Consistently, the other tools displayed lack of power in the real LCA and RP cohorts with mixed ethnicity and high locus heterogeneity.

The performance advantage of GRIPT might be partly due to that it scores the mutation load of a gene according to the Mendelian inheritance rule. Under the AR model, for each individual, GRIPT only considers/scores genes with at least two variants, which could exclude the false positive signals from the genes merely carrying one pathogenic allele in an individual. Furthermore, the Fisher’s test built upon the combination of a binomial test and a WRS test equipped GRIPT the excellent statistical power for comparing highly skewed distributions of gene score (Figure 1 and Methods). In comparison, VAAST/VAAST2, CMC, SKAT and KBAC takes into account the genes carrying at least one variant in an individual. In addition, CMC, SKAT, and KBAC group all the variants within a gene to compute the deleterious mutation load of the gene, which makes genes with large number of rare variants in case cohort (e.g. benign or due to chance) ranked high and creates false positives. As shown in simulation, this impact on the other tools was more pronounced when the true signal was diluted by high locus heterogeneity and/or was compromised by large background noises, e.g. population stratification (or sequencing platform/variant calling difference) or relaxed cutoff of variant filtering frequency.

Simulation results also suggest that to optimize the performance of GRIPT, the following conditions should be considered. First, as one of the key factors affecting sensitivity is the proportion of patients attributed to the same gene, it is highly desirable to increase the homogeneity of patient cohort. One possible approach is to perform detailed phenotyping and gather the patients who share similar phenotypes and are likely due to mutations in one or a small number of genes. Second, while maintaining the homogeneity of the patient cohort, increasing the patient cohort size can also improve sensitivity. For example, by increasing the patient cohort size from 50 to 100 while maintaining 2% of patients carrying disease mutations of the same gene under the AR model, the average rank of the disease gene increased from 31 to 3 by GRIPT. Third, using the correct inheritance model when running GRIPT can leverage its power. If the inheritance model of the diseases is unclear, GRIPT should be run using different models, including AD, AR, XD and XR, respectively. Fourth, reduction of the noises in the input variants will improve the outcome. For example, large databases of “normal” populations, e.g. gnomAD and ExAC should be used to pre-filter variants and remove common benign variants that are unlikely to cause diseases, while filtering with internal databases can weaken the error/bias from the sequencing platforms and variant callers. Furthermore, under different inheritance models, the mutations should be pre-filtered with different frequency cutoffs (for example, the variant filtering frequency for AD model should be more stringent, namely lower than for AR model). Additionally, removing the genes that are highly mutable but known not causing diseases can reduce noise as well. Fifth, the accuracy of variant function/pathogenicity prediction will also impact the performance of GRIPT. Currently GRIPT applies the well-established integrative allele prediction score, i.e. CADD score, to predict the pathogenicity of variants. However, as the scoring system of GRIPT is flexible, users can easily substitute the CADD score with any other score generated by better algorithms for variant pathogenicity prediction. In aforementioned analysis, we also used DANN and REVEL scores as the variant score, which generates the consistent results, suggesting the reliability and robustness of the statistic test framework of GRIPT. The thumb of rules for using variant score systems is: 1) the scoring systems should reliably and quantitatively predict the deleteriousness of variants. 2) the scores should be scaled/normalized into a genome-wide ranked score to allow the comparison implemented in the statistic test of GRIPT. 3) The score system should be comprehensive and cover all the possible SNP and INDEL in the genome.

Although GRIPT does not directly identify pathogenic mutations, by identifying candidate (novel) disease genes, it will dramatically reduce the number of variants to be considered for each patient and therefore greatly facilitate the identification of potential mutations. Once the candidate genes are identified, the causal variants of the genes can be further prioritized with the conventional steps: 1) The individuals carrying at least two (recessive mode) or one (dominant mode) rare variants of the candidate gene should be identified from the patient cohort. 2) Multiple variant effect prediction systems can be applied to estimate and compare deleteriousness of the variants in affecting protein function, mRNA splicing or other regulation processes of the gene (e.g. CADD score, SIFT, Polyphen, MetaLR/SVM, PROVEAN, REVEL, phyloP100way_vertebrate, phastCons100way_vertebrate, ada_score, NNsplice). 3) The sanger validation and segregation tests of the patients and additional relatives should be performed for the candidate variants.

### Conclusions

In summary, we developed a highly accurate and robust case-control analysis method, GRIPT, for discovery of Mendelian disease genes. It is especially powerful in detecting disease genes underlying diseases with high locus heterogeneity and is less affected by population stratification. It is also efficient, portable, and flexible. In addition, we generated a WES data simulator which is capable of unbiasedly simulating the WES data of control cohorts with any sample size, gender ratio, and population ethnicity for the usage of GRIPT or other tools. As NGS technology advances (e.g. the decrease in cost and time) and greater amounts of large cohort data become available, we envision that GRIPT will make a significant contribution to the discovery of novel Mendelian disease genes and pave the way for better understanding, diagnosis, prevention, and treatment of Mendelian diseases.

## Methods

### Each variant is scored to quantify the deleteriousness

The hypothesis that GRIPT tests is whether the deleterious mutation loads of a disease-causing gene is significantly higher in case cohort than in control cohort. To quantify the deleteriousness of variants, in this study, we applied Combined Annotation Dependent Depletion (CADD v1.3) score to each variant of each gene in every individual [32]. CADD score is an integrative score derived from the integration of diverse annotations and is highly predictive of molecular functionality and pathogenicity [32]. Higher CADD score indicates more deleteriousness of the mutation. In addition, CADD not only provides integrative prediction scores for SNVs but also for INDELs which are missing for most other variant effect prediction tools. We further normalized the variant score on a scale of 0 to 1 as *s* = 1 − 10^−*C*/10^. *C* is the PHRED-like scaled C-score as described in CADD. Moreover, CADD score can be easily replaced by any other score that users provide in order to better predict the variant’s deleteriousness. To test the reliability and robustness of the statistic test framework of GRIPT, the ranked REVEL and DANN scores were also applied as the variant scores respectively. The CADD score was downloaded from https://cadd.gs.washington.edu/download. The ranked DANN score was extracted from dbNSFP3.4a downloaded from https://sites.google.com/site/jpopgen/dbNSFP. The ranked REVEL score was downloaded from https://sites.google.com/site/revelgenomics/downloads.

### Each gene is scored under different inheritance models

Under the autosomal recessive (AR) model, only the genes with at least two variants in an individual will be assigned a positive score. The sum of the two highest scores of variants within a gene is used as the score of that gene in the individual. If two variants of a gene are *in cis* (namely, the two variants reside on the same chromosome) in an individual, only the variant with the higher score will be considered. If a gene carries ≤ one variant in an individual, the score of this gene will be 0 for that individual. Under the AR model, the maximum score for a gene is 2, and the minimum is 0.

Under the autosomal dominant (AD) model, only the genes with at least one variant in an individual will be assigned a positive score. The highest score of variants within a gene is used as the score of that gene in the individual. Under the AD model, the maximum score for a gene is 1, and the minimum is 0.

Similarly, under the X-linked recessive model, the sum of the two highest variant-scores is used as the score of each gene on the X chromosome in an individual. And under X-linked dominant model, the highest variant-score is used as the score of each gene on the X chromosome in an individual.

### Gene score distribution is highly skewed for rare Mendelian disorders

As mentioned above, each gene has a score in each case or control individual, ranging from 0 to 1 (for dominant models) or 2 (for recessive models). Then, for each gene, we compare the gene score distribution in case cohort to that in control cohort. The null hypothesis is that the deleterious mutations load of a gene is not significantly different between cases and controls. Thus, the significance of the one-tailed alternative hypothesis that the deleterious mutations load is higher in cases than controls could suggest the likelihood of the gene associated with the disease.

To choose the appropriate statistic test, we first characterized the gene score distribution. We found that the score distributions of most genes are highly skewed with excesses of zeros. This is expected mainly because Mendelian diseases are rare and so are the disease-causing mutations. Usually, after filtering out known common human variants which are likely benign, only a small number of rare variants (e.g. MAF ≤ 0.5%) in cases and controls will be kept. Moreover, among the filtered rare variants, only some of them have deleterious effects, therefore, only these rare, deleterious variants will have positive variant-scores. In addition, the recessive model requires a biallelic state to assign a positive gene score in one individual. Thus, the scores of a gene in most case individuals and control individuals are zeros. An example of *USH2A* gene score distributions in our retinal disease patient cohort (n = 250) and an internal control cohort (n = 250) is shown in Figure 2.

### Combining two separate statistical tests with Fisher’s test

To compare the highly skewed distributions of gene scores in case and control cohorts derived above, we test a composite null hypothesis by applying a Fisher’s test to combine two separate tests including a binomial test and a WRS test [19]. The composite null hypothesis is designed to answer two questions. The first question is whether the proportions of non-zero scores are similar in case cohort and control cohort (Z_1_ =0). The second question is whether the values of non-zero scores are similar in case cohort and control cohort (Z_2_ = 0). Namely, Fisher’s method will test the H_0_: Z_1_ =0 and Z_2_ =0 versus the one-tailed alternative H_1_: Z_1_ > 0 and/or Z_2_ >0[19].

Let *N*_1_ and *N*_2_ be the total number of cases and controls. Let *n*_1_ and *n*_2_ be the number of non-zero score in cases and controls respectively.

The first statistic, *Z*_1_, represents the proportion difference of non-zero scores between cases and controls. Given *n*_1_ + *n*_2_ = *n* and *r* = *N*_2_/*N*_1_, *n*_1_ is approximately distributed as *Binomial*(*n*, 1 + *r*)^−1^) under H_0_. Hence, a one-tailed p-value *p*_1_ can be obtained as the tail area under the *N* (0, 1) p.d.f to the right of

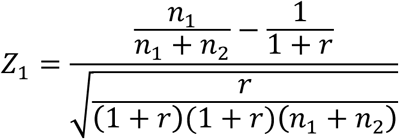

The second statistic, *Z_2_*, represents the difference of the non-zero scores between cases and controls. The standardized Wilcoxon rank sum test was applied to test whether the gene cores in cases are significantly higher than those in controls. Let *p*_2_ enote the corresponding one-tailed p-value.

Finally, Fisher’s method is used to test the composite null hypothesis H_0_: Z_1_ =0 and Z_2_ =0 at one-tailed level *α* based on a combination of Z_1_ and Z_2_ or *p*_1_ and *p*_2_ as follow:

Reject H_0_ if *p* < *α*, where 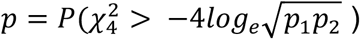

Here, 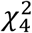 is a χ^2^ distribution with 4 d.f., therefore the p value can be calculated using a χ^2^ distribution.

The program of GRIPT is written in Java and R.

### A WES data simulator based on ExAC database

The VCF file of ExAC database (ExAC.r0.3.1.sites.vep.vcf) was downloaded from http://exac.broadinstitute.org/downloads [33]. We collected the variants recorded in the VCF file which were not indicated as filtered by ExAC. For each of these variants, we extracted information on the genomic position, the allele count, the chromosome number, and the allele frequency in each subpopulation, including AFR (African/African American), AMR (American), EAS (East Asian), FIN(Finnish), NFE (Non-Finnish European), SAS (South Asian), OTH (Other), adjusted population, and raw data. We only considered the ExAC variants that were missense or loss-of-function mutations (e.g. missense mutations, stop-gained mutations, splicing donor/acceptor mutations, and frameshift mutations). We also downloaded the CADD scores for the ExAC variants from http://cadd.gs.washington.edu/download [32] and annotated each collected ExAC variant with its corresponding CADD score. Next, we wrote a WES data simulator program in PERL. Briefly, the script simulated the WES data per person individually. For each individual, the simulator will go through the variants recorded in the ExAC database which satisfy the variant filtering criteria (e.g. MAF ≤ 0.5%) one by one and output the reference nucleotide or the altered nucleotide according to the allele frequency of that variant in ExAC. For example, in the position chr1:10000, if the allele frequency of “A>T” is 0.2% and the allele frequency of “A>G” is 0.5%, then in the simulated WES data of one person, there is 0.2% of chances the simulator will output the SNP “A>T”, 0.5% of chances will output the SNP “A>G”, and 99.3% of chances the simulator will generate “A>A”, namely not output any SNP in chr1:10000. Thus, each generated variant follows a multinomial distribution according to its frequency in the user-selected ethnic population based on the ExAC database. For a given number (N) of individuals with a given sex ratio, the simulator will generate “N” WES data file individually. Each WES data file includes information such as reference nucleotides, altered nucleotides, the coordinates in the genome, and the CADD scores of the variants.

### Simulation of patient and control cohorts

To evaluate the performance of GRIPT, we performed the simulation tests on GRIPT and similar tools, i.e. VAAST2, CMC, SKAT and KBAC. The WES data of the patient cohort and control cohort were first generated using the WES data simulator mentioned above. Given the rare frequency of Mendelian disease-causing variants in normal population, for the AR model, the WES data were simulated based on the variants whose maximum population frequency was ≤ 0.5% in ExAC database by default, while for the AD model, based on the variants whose maximum population frequency was ≤ 0.01% in ExAC database by default, unless otherwise specified. We used “adjusted” average population frequency as the default variant frequency, unless otherwise specified. Then, we randomly selected pathogenic variants of a given disease gene from HGMD database with MAF ≤ 0.5% in ExAC database, and inserted them into a given percentage of individuals randomly selected from the patient cohort to mimic the patient cohort with genetic heterogeneity. In the AR model, two variants were respectively selected from HGMD and spiked into each selected individual. Thus, the two variants spiked into the same individual can be the same (homozygous) or different (heterozygous). In the AD model, only one variant was randomly selected and spiked into each selected individual. Therefore, under the AR or AD model, the pathogenic mutations of a given gene can be the same or different within and between the patients. No additional mutations were spiked into the control cohort. For each spike-in percentage level per scenario, 30 simulation runs were repeated (Supplementary figure S1).

### The implementation of VAAST2, CMC, SKAT, KBAC and Fisher’s exact test

The latest release of VAAST2 was obtained from http://www.yandell-lab.org/software/vaast.html [16, 17]. The CMC, SKAT and KBAC were implemented through the “Rvtests” software package downloaded from https://genome.sph.umich.edu/wiki/Rvtests#Download [34]. The p-values of VAAST2, SKAT, and KBAC were obtained using 400000 permutations. The Fisher’s exact test was implemented through the PLINK v1.90b5.2 package from https://www.cog-genomics.org/plink/1.9/ [35]. The intermediate steps were carried out using PERL and R scripts.

### Preprocessing the variants in cis

To reduce false positive, we recommend the users to handle the variants *in cis* before inputting data into GRIPT. However, given that it is not always possible to obtain accurate phasing information, GRIPT can tolerate imperfect phasing as shown in the aforementioned simulation and real data analyses. Currently, a preprocessing script included in the GRIPT package was used to handle variants *in cis*, which perform the following operations:

1) If the genomic coordinates of two variants are within 100bp, Fisher’s exact test will be performed to determine whether the two variants are *in cis* by comparing the ratio of the variant base sequencing coverage to the reference base sequencing coverage of the two variants. If the two variants are *in cis* and within 100bp, they can be covered by a large number of the same sequencing reads, therefore their read coverage ratios would be similar and Fisher’s exact test p-value would be large. In contrast, if they are *in trans* and close to each other, they would be covered by different sequencing reads, thus the read coverage ratios of the two variants would be different and Fisher’s exact test p-value would be small. We take Fisher’s exact test p ≥ 0.4 as the cutoff to deduce the read coverage ratios of the two variants are similar, namely, the two variants as *in cis*, otherwise as *in trans*. Using different p-value cutoff does not significantly impact on the result. For example, we have used p < 0.05 as the cutoff to assign the variants *in trans*, and p ≥ 0.05 to assign the two variants *in cis*. Although this could mistakenly assign a few *in-trans* variants as *in-cis*, the results remained consistent. Because GRIPT is built on the mutation burden in case cohort and control cohort but not a single case, a few imperfect phasing cases can be tolerated. If the two variants are determined to be *in cis* by Fisher test, the variant with higher variant score (e.g. CADD score) will be passed on to the subsequent analysis, while the one with lower variant score will be ignored.

2) For each gene in every individual, all variants within the gene will be searched against the same gene in the rest individuals of the case cohort. If a gene has ≥ 2 variants present concurrently in ≥ 2 individuals, it is likely that these variants are *in cis*. Because given the sample size of case cohorts (n = 115 for LCA, 154 for the RP cohort, and currently available case cohort size mostly ≤ 5000) and the rare frequency of Mendelian disease-causing mutations (allele frequency ≤ 0.5%), the chance for two or more rare variants co-occurring in unrelated individuals is very small (5000 * (0.005*0.005)^^2^ << 2), unless these variants are *in cis* or the disease is specifically caused by the combination of the variants. Although our preprocessing script does not fit the latter situation, it can help clean up the former one. If a gene has ≥ 2 variants co-occurring in ≥ 2 individuals, among the concurrent variants, the script will only keep the variant with highest variant score and ignore the other concurrent variants in the subsequent analysis.

## List of abbreviations

AFR: African/African American
AMR: American
AR: Autosomal recessive
AD: Autosomal dominant
CADD: Combined Annotation Dependent Depletion
CAST: Cohort Allelic Sums Test
CMC: Combined Multivariate and Collapsing
DANN: Deleterious Annotation of genetic variants using Neural Networks
EAS: East Asian
ExAC: Exome Aggregation Consortium
FIN: Finnish
gnomAD: genome Aggregation Database
GRIPT: Gene Ranking, Identification and Prediction Tool
GWSL: Genome-wide significant level
HGMD: Human Gene Mutation Database
KBAC: Kernel-Based Adaptive Clustering
LCA: Leber’s congenital amaurosis
NFE: Non-Finnish European
NGS: Next generation sequencing
NSAG: Number of significant autosomal genes
OMIM: Online Mendelian Inheritance in Man
OTH: Other
REVEL: Rare Exome Variant Ensemble Learner
RP: Retinitis pigmentosa
SAS: South Asian
SKAT: Sequence Kernel Association Test
VAAST: Variant Annotation, Analysis and Search Tool
VCF: Variant Call Format
WES: Whole exome sequencing
WGS: Whole genome sequencing
WRST: Wilcox rank sum test

## Declarations

### Ethics approval and consent to participate

All subjects in the study underwent ophthalmic evaluations. Informed consent was obtained from all patients or their guardians. All the diagnostic procedures were approved by the local institutional review boards or ethics committees.

### Availability of data and materials

The GRIPT software, scripts, WES data simulator, and test examples can be downloaded from github: https://github.com/fe4960/GRIPT_BCM And zenodo: https://zenodo.org/record/1407225#.W4msEdhKgb0 DOI:10.5281/zenodo.1407225

The simulated datasets can be downloaded from: ftp://ftp.hgsc.bcm.edu/RChen/JWang/Simulation_data.tar.gz

## Authors’ contribution

J Wang, L Zhao, X Wang, F Wang, R Chen proposed the method. J Wang, L Zhao, X Wang wrote the software. J Wang, L Zhao, Y Chen performed the simulation test. J Wang, M Xu, Z Ge, Z Soens analyzed the patient data. J Wang, L Zhao, X Wang, P Wang, R Chen wrote the manuscript.

## Supplementary Figures (.pdf format)

Supplementary Figure S1: The main procedure of simulation analysis

Supplementary Figure S2: Benchmark of GRIPT with REVEL and DANN scores on 400 Mendelian disease genes.

Supplementary Figure S3: Test the impact of patient cohort sizes with REVEL and DANN scores

Supplementary Figure S4: Test the impact of population stratification with REVEL and DANN scores

Supplementary Figure S5: Test the impact of variant frequency filtering with REVEL and DANN scores

## Supplementary Tables (.xls format)

Supplementary Table S1: The sensitivity and specificity of GRIPT and other tests under the AR and AD models

Supplementary Table S2: Benchmark on 400 randomly selected known disease genes

Supplementary Table S3: Test the effect of the patient cohort sample size

Supplementary Table S4: Test the effect of Population stratification in cohorts

Supplementary Table S5: Test the effect of variant frequency filtering

Supplementary Table S6: Test the effect of the control cohort size

